# Combinatorial CRISPR/Cas12a excises targeted loci in the mammalian genome at very low efficiency

**DOI:** 10.64898/2026.09.21.753273

**Authors:** Nazanin Esmaeili Anvar, Zsofia M Szegletes, Ryan J Steger, Abby V McGee, Sidney Wang, John G Doench, Traver Hart

## Abstract

The CRISPR/Cas12a system has emerged as a highly useful tool for genomic editing due to its ability to process multiple guide RNAs (gRNAs) from a single promoter. In this study, we explored the potential of combinatorial CRISPR/Cas12a to excise targeted loci in the mammalian genome by inducing double-strand breaks (DSBs) upstream and downstream of targeted exons. We designed a library of gRNA pairs targeting introns flanking 54 exons of 18 essential genes and 17 exons of 10 nonessential genes, aiming to assess the efficiency of Cas12 in excising targeted loci. While single gRNAs directly targeting essential exons produced expected fitness defects in cells, pairs of guides targeting flanking intronic regions resulted in only trivial viability effects, suggesting that excision rarely occurred. Further analysis showed no meaningful correlation between gRNA efficiency, orientation, GC content, or cut site geometry and the success of locus excision. These findings indicate that, under the conditions tested, the Cas12a system does not yield locus excision at the efficiency required for sensitive negative selection assays.

## Background

Less than 2% of the human genome codes for proteins, while the vast majority consists of noncoding sequences that may still play critical roles in cellular function^1^. Notably, about 93% of disease- and trait-associated variants are found in these noncoding regions^2^, highlighting their potential impact on human health and the need to better understand their functions. There has therefore been considerable interest in CRISPR-mediated genetic engineering systems to characterize these elements by enabling efficient disruption of noncoding elements within their natural chromatin environment in a high-throughput manner^3^. While the study of noncoding regions has been conducted in various pooled CRISPR studies using different methods, it is important to note that when targeting less understood genetic sites or non-coding regions, the density of available PAM sequences^4^ and the presence of highly efficient gRNAs may be less frequent. Historically, the main application of Cas9/Cas12a is to introduce double-strand breaks and error-prone repair resulting in indels, which can introduce frameshifts and loss of function (LOF) in protein coding genes but may not disrupt function of noncoding regions.

A better way to understand the functional relevance of noncoding DNA involves physically removing specific loci from the genome and observing the subsequent changes in cell phenotype. To excise targeted genomic loci in cells, CRISPR technology can in principle be employed to create precise cuts upstream and downstream of the locus. Previous studies have attempted this, employing Cas9 to delete noncoding regions in the genome ^5–7^. For example; Aparicio-Prat *et al*. designed an experiment to delete the promoters of one protein-coding gene and two oncogenic long noncoding RNAs ^6^. Similarly, Diao *et al*. examined a 2-Mb region near the POU5F1 gene locus. They employed 11,570 pairs of sgRNAs in human embryonic stem cells and reported the detection of 45 cis-regulatory elements^5^.

While the Cas9 enzyme derived from Streptococcus pyogenes (SpCas9) remains the preferred choice for a wide range of genetic screening studies, the effective delivery of multiple guide RNAs for pooled screening has posed challenges due to the requirement for individual promoters for each single guide RNA ^8^. In contrast, the Cas12a CRISPR endonuclease allows the expression of an array of guide RNAs from a single promoter that the Cas12a enzyme can process into independent targeting units ^8,9^. These distinctions in processing mechanisms and the ability to express multiple gRNAs into a single construct could significantly simplify the process of designing and delivering combinatorial genome editing tools.

We previously compared the efficiency of Cas9 and Cas12 in multiplexed knockout CRISPR studies by conducting a meta-analysis of studies that applied different methods to assess genetic interactions between gene pairs using dual knockout arrays. Parrish ^10^ and Thompson ^11^ utilized SpCas9 with two gRNA expression cassettes, while Ito ^12^ used orthogonal Cas9 proteins from *Staphylococcus aureus* and *Streptococcus pyogenes*. Gonatopolous ^13^ implemented a hybrid Cas9/Cas12a approach (CHyMErA), using the endogenous RNA processing capability of Cas12a to cleave a guide array into independent Cas9 and Cas12a guides. We used the engineered enAsCas12a system, an efficiency-optimized Cas12a enzyme^8,14^, and showed multiplexing capability of enCas12a when compared to Cas9 ^15^. This advancement showcases enCas12a’s improved ability to simultaneously target and perturb multiple genetic loci ^15,16^.

In this study, we put forward the idea that the inherent multiplexing capability of enCas12a could be leveraged to simultaneously introduce two cleavage sites, one upstream and one downstream of the intended region. This approach would lead to the excision of the desired genetic segment, allowing for phenotypic assays to discover locus function. By doing so, we can not only study the effects of non-coding regions, enhancers, and promoters but also identify the function-specific exons within protein-coding genes. Ultimately, we found that the system lacked the efficiency required for effective use at scale.

## Results and discussion

Our systematic investigation aimed to identify the key factors that contribute to the effectiveness of gRNA pairs in achieving precise DNA excision. In this experiment, we attempted to excise exons in known essential and nonessential genes to assess enCas12a’s ability to remove genomic loci, with the expectation being that excision of a constitutive exon of an essential gene would yield a fitness phenotype similar to direct targeting of the exon (i.e. gene knockout). A key advantage of the Cas12a system is that it can process an array of guide RNA sequences expressed from a single Pol III promoter into a set of independent, active guides, with no obvious positional effects^15–17^. We reasoned that, by simultaneously targeting introns both upstream (5’) and downstream (3’) of essential exons, we could readily identify “essential” guide pairs, and by doing so at scale we could learn the sequence and target features that maximize the efficiency of this assay.

We selected genes from established reference sets of essential and nonessential genes^18,19^. We selected 54 exons in 18 essential genes and 17 exons in 10 nonessential genes from the Hart reference gene sets, and tested both exonic (gRNAs targeting exons) and intronic gRNAs. Each exon’s upstream and downstream intron was targeted by ∼30 different intronic gRNAs (∼900 unique pairs) with different criteria (Fig. 1A-B). As controls, we used guides directly targeting the exons of the 18 essential and 10 nonessential genes, paired with guides targeting known nonessential genes (Fig 1B). We additionally used single guides targeting a broader selection of exons within the same 18 essential (88 exons) and 10 nonessential (41 exons) genes, to measure phenotypic consistency of the gene knockout across exons. Total library size is 77k unique guide pairs, with 46k guide pairs targeting introns bracketing essential exons, 12k guide pairs with a single intronic guide coupled with a guide targeting a nonessential gene, and 15k guide pairs targeting introns bracketing nonessential exons.

**Figure 1.**
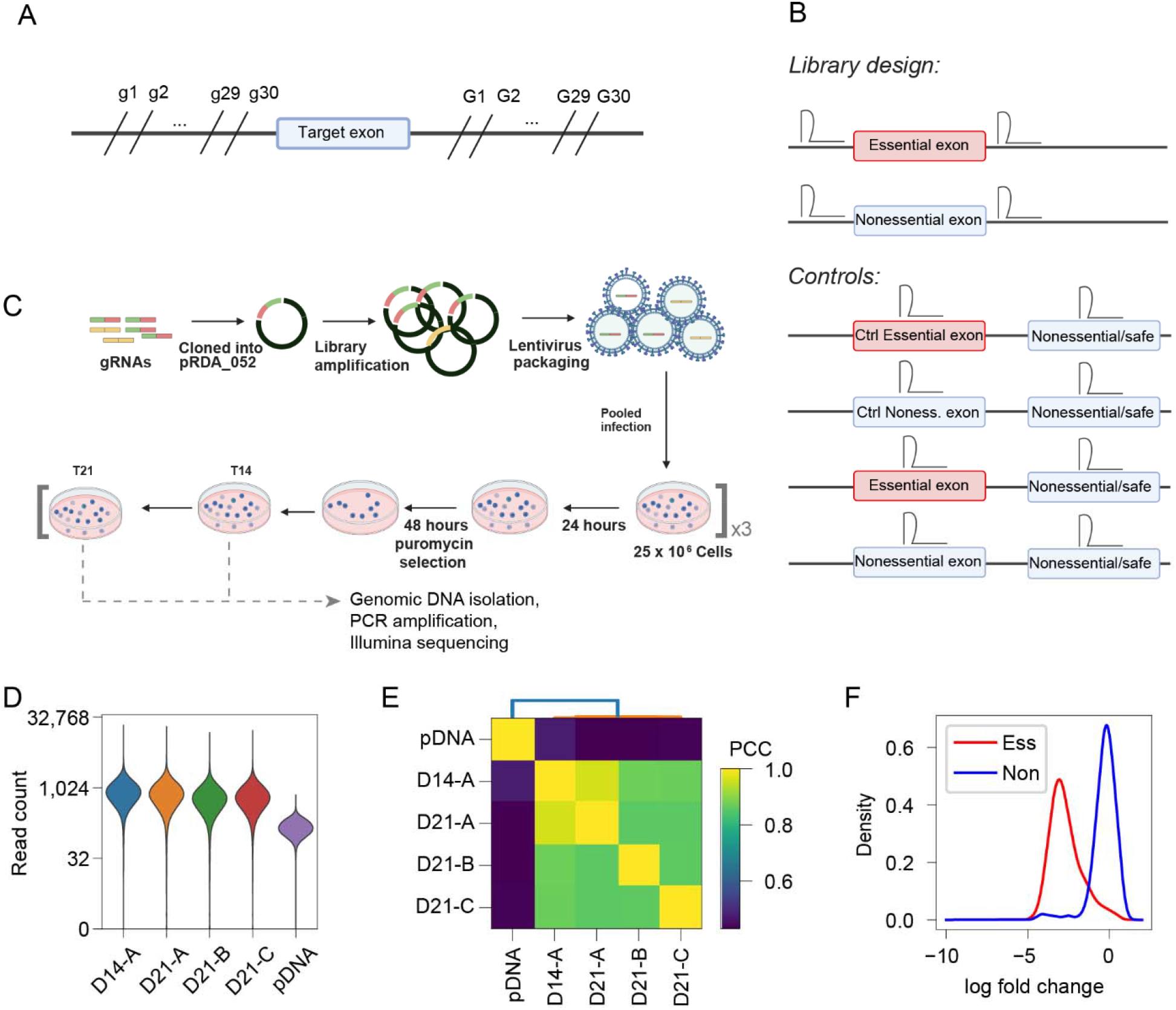
Experimental design and QC. (A) Each experimental exon was targeted by 900 pairs of intronic constructs, comprised of all pairs of 30 gRNAs upstream and 30 gRNAs downstream of a target exon. (B) Library constructs. Experimental group targets introns up and downstream of target exons singly (not shown) and in pairs. Controls target exons of reference essential and nonessential genes, plus target exons of intronic constructs, plus additional exons (‘broad’) of these genes, to confirm target expected behavior when exon is excised. Each exonic guide RNA was paired with a corresponding negative control targeting a reference nonessential gene. C) Experimental workflow to screen the library in A375 cell line. (D) Quality control: read count distribution for sequenced samples. (E) Quality control: correlation of read counts by sample. (F) Quality control: guides targeting reference essentials (red) vs. guides targeting reference nonessentials (blue), mean of D21 samples.

The library was screened in A375 cells using standard protocols. Oligos containing the guide pairs were commercially synthesized and cloned into the pRDA_052 guide expression vector, sequence-verified, packaged in lentivirius, and transduced into A375 melanoma cells. Transduction and passaging were performed at scale to maintain 500x library coverage, and cells were collected and sequenced at three timepoints T14 and T21 (Fig. 1C). The screen passed all quality control checks: both control (plasmid) and experimental (endpoint) samples yielded good readcounts, and the library was well represented in all samples (Fig 1D). Guide array readcount profiles were highly correlated across samples (Fig. 1E), and exonic guides targeting essential genes were strongly separated from guides targeting nonessential genes (Fig. 1F). Collectively, these measures are indicators of a successful pooled library gene knockout experiment.

To evaluate whether the system effectively excised targeted loci, we considered the fold change distributions of guide arrays in the library. Arrays containing a gRNA directly targeting exons of essential genes showed an expected strong negative fold change (Fig. 2A, Exonic-ess:Neg_controls), compared with minimal fold change of arrays containing gRNA only targeting exons of nonessential genes (Fig. 2A, Exonic-noness:Neg_controls). These results from positive and negative controls confirm the on-target cutting ability of our constructs using enCas12a, as well as the cell-line specific phenotype of the targeted genes. However, guide arrays targeting upstream and downstream introns of essential exons (Fig. 2A, Intronic-ess:Intronic-ess) showed only a minor shift relative to guide arrays targeting upstream and downstream introns of nonessential exons (Fig. 2A, Intronic-noness:Intronic-noness). Though this shift is statistically significant (P-value < 2.2×10^-16^, independent t-test), the P-value is due to the large sample size (46,431 intronic clones targeting essential genes vs 15,062 intronic clones targeting nonessential genes), but the effect size is not large and the observed fold change is still near zero. In fact, the fold change distribution of arrays designed for essential exon excision was almost identical to the fold change distribution of arrays with one essential intron guide paired with a negative control guide (P-value = 2.91×10^-14^, independent t-test, n= 12,465 clones targeting Exonic-ess paired with negative controls, but Cohen’s d = 0.08; Fig. 2B). In summary, a positive outcome of this experiment would be for exon excision to phenocopy direct exon targeting, but instead the fold change distribution of exon excision constructs almost exactly mirrors that of constructs single targeting the introns of essential genes, indicating very low efficiency of locus excision.

**Figure 2.**
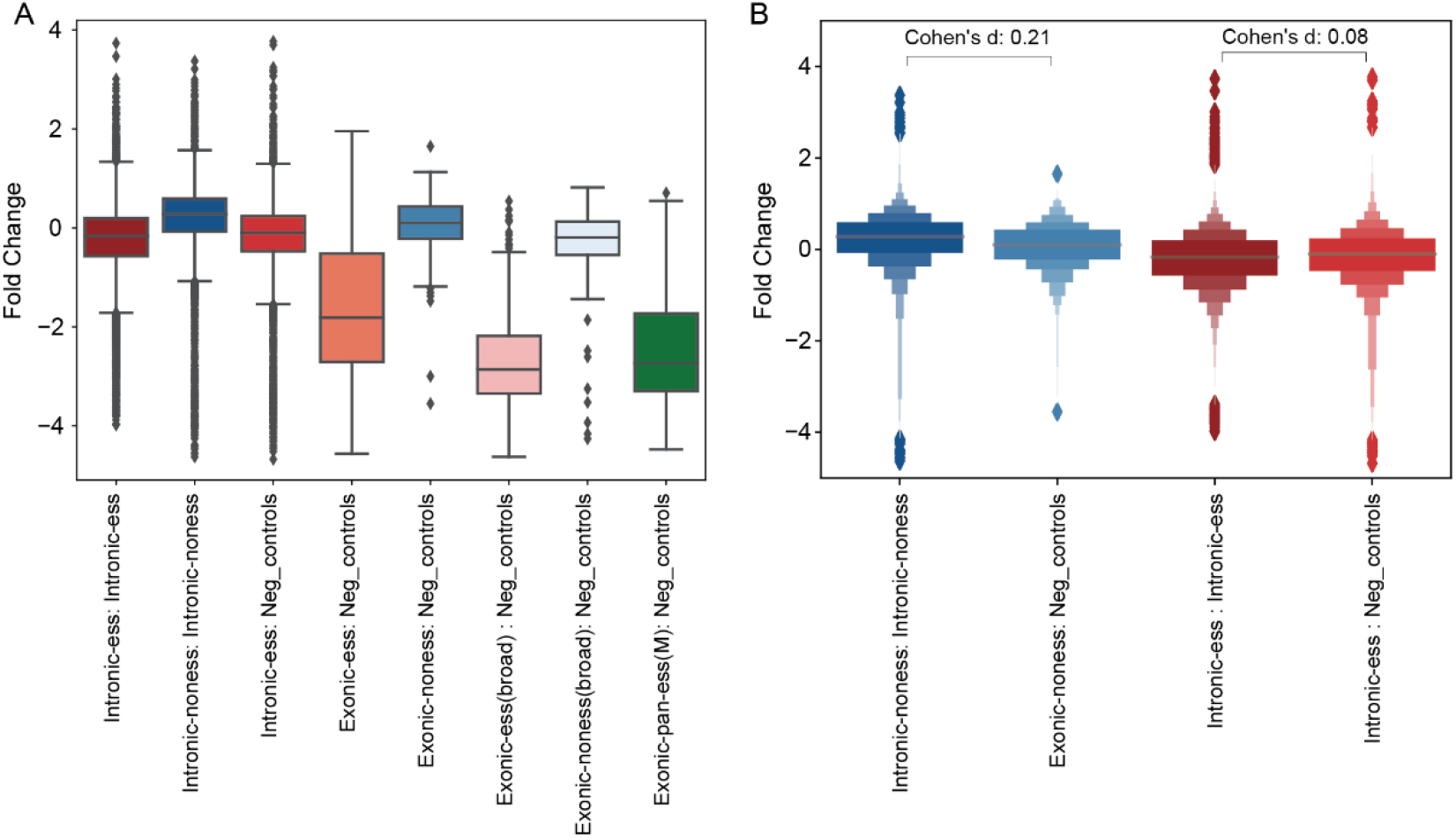
Fold change by target class. A) Fold change distributions across all target classes. Combinatorial targeting of introns 5’ and 3’ of essential exons (Intrinic-ess:Intronic-ess) did not show signature consistent with essential gene knockout, while single guides targeting essential exons (Exonic-ess) showed expected fold change. B) Comparing similar target classes. Blue, single-targeting nonessential exons (exonic-noness) vs. dual-targeting introns flanking nonessential exons. Red, single-targeting vs. dual-targeting introns flanking essential exons.

Though the overall experiment did not appear to work as we had hoped, the distribution of fold changes for guide arrays targeting essential exon excision showed a long tail of negative fold changes. Therefore, we explored the resulting data to attempt to identify factors that could improve performance of locus excision and to determine whether any features associated with guide array design could bias results toward more successful outcomes. We examined efficiency of gRNAs, GC content, distance from cut sites, orientation of gRNAs and homology sequences around the cuts for whether these factors could bias the results toward successful outcomes.

First, we considered the efficiency of individual guides in our positive and negative controls. As all guides were designed using CRISPick, we compared the knockout phenotype of single guides directly targeting essential exons with the predicted on-target efficacy score (Fig. 3A). Efficacy score is strongly predictive of knockout phenotype. Fold change distribution of 1,046 positive control arrays (exonic gRNAs targeting essential genes) had a significant correlation between gRNA efficiency and screen dropout (R^2^ = 0.22), and guides with on-target efficacy scores >= 0.7 consistently showed strong fold change when targeting essential genes. This result confirms CRISPick’s prediction scores for ranking efficient gRNAs. The on-target efficacy scores of exonic guides targeting nonessential genes (330 arrays) did not show a strong correlation, as expected (Fig. 3B). We also considered the GC content of gRNAs as a factor that could influence screen efficiency, but no correlation was observed between gRNA GC content and fold change of guides targeting reference essential (Fig 3C) or nonessential (Fig 3D) genes.

**Figure 3.**
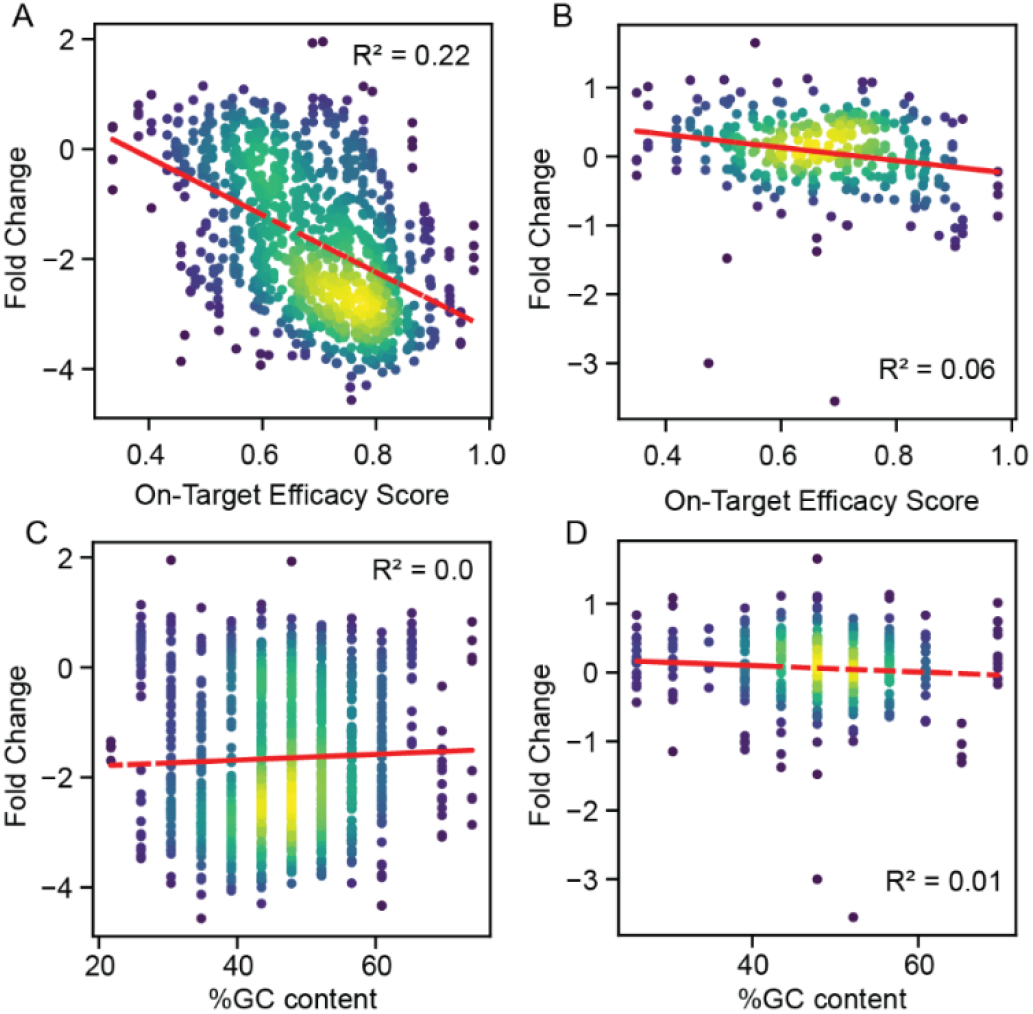
On-target efficacy score is a good predictor of guide efficacy. (A) For all guides targeting exons of essential genes, the on-target efficacy score is plotted against observed fold change, colored by relative density. (B) Same plot for guides targeting exons of nonessential genes. (C-D) gRNA GC content vs fold change of guides targeting exons of essential (C) or nonessential (D) genes.

Next, we investigated whether the same correlation observed between on-target efficacy scores and array fold change in exonic guides could also be identified in intronic guides. The data from intronic gRNA pairs showed that higher efficiency of each of the gRNAs of a pair did not meaningfully correlate with a greater fold change (R^2^ = 0.04 and R^2^ = 0.02 for guide 1 and guide 2, respectively) (Fig. 4A-B). This suggests that higher guide efficacy did not enhance the excision efficiency of the targeted loci. The selection process for intronic guides required that the CRISPick on-target prediction score > 0.6, which is in the effective region of Fig 3C, so most of these guides should be effective and variation of guide score within this narrow range may not be predictive of better performance.

**Figure 4.**
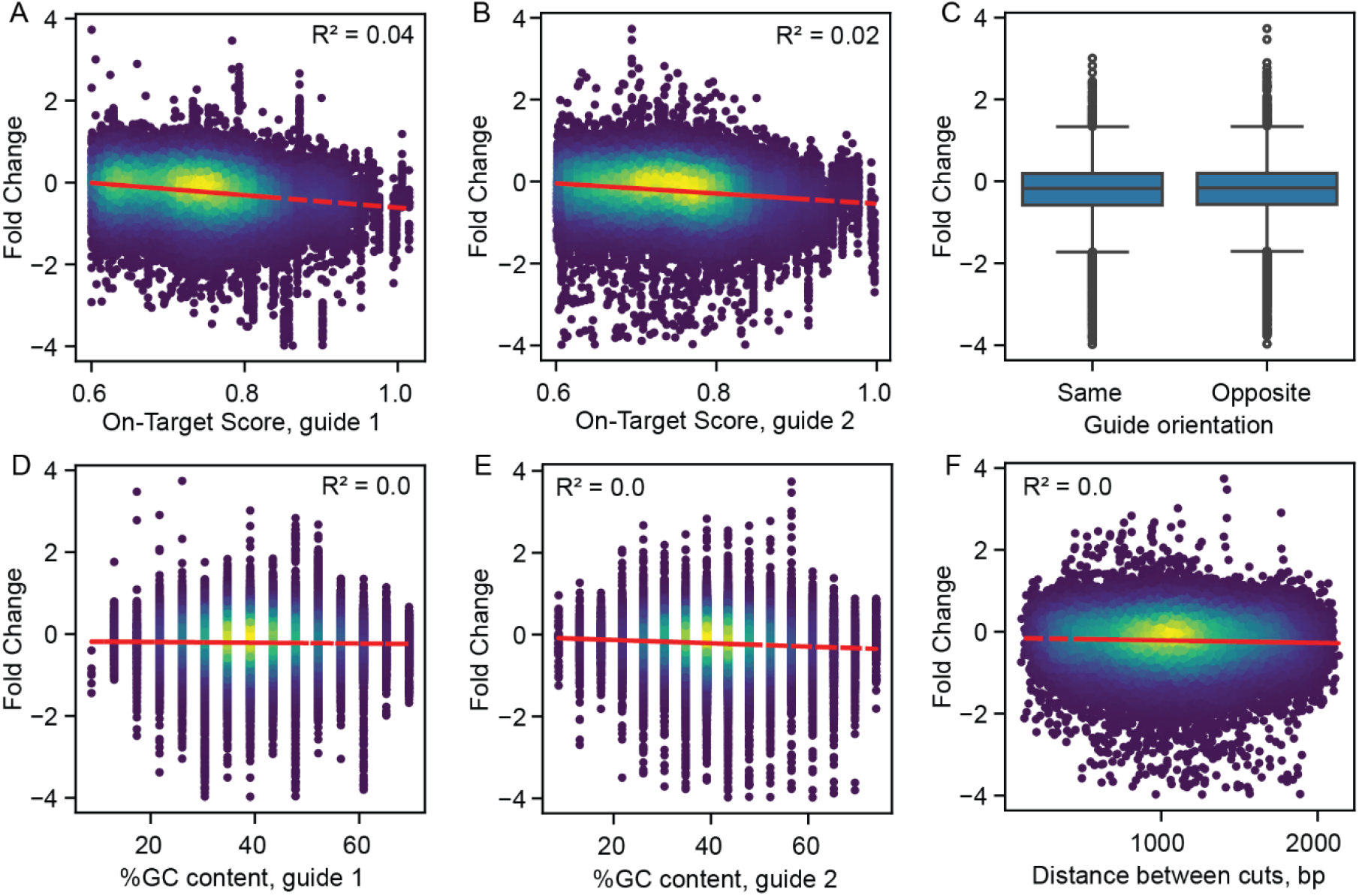
Guide and target characteristics do not optimize excision. A) On-target score from CRISPick vs observed fold change, for guide 1 in guide pairs targeting introns of essential exons. (B) Same plot for guide 2 of the guide pairs. (C) Fold change of guide pairs targeting introns of essential exons, where target sites are in the same or opposite orientation on the genome. (D-E) GC content of guide 1 (D) and guide 2 (E ) of intronic gRNAs. (F) Distance between cut sites vs. fold change, for guide pairs targeting introns of essential exons.

Every set of intronic gRNAs could consist of two gRNAs with target sites oriented in the same direction or in opposite directions. To investigate whether one of these orientations yielded more favorable outcomes in terms of excision efficiency, we systematically classified these gRNA pairs into two categories: “Same Orientation” and “Opposite Orientation.” Neither group exhibited an advantage in terms of enhancing the excision efficiency (Cohen’s D = 0.01: Fig. 4C). Similar to exonic gRNAs, where it was shown that the cutting efficiency is independent of the gRNA GC content, intronic gRNAs also did not demonstrate improved efficiency when the GC content percentage was varied. (Fig. 4D-E).

Next, we considered the distance between the two cut sites. Inducing two simultaneous Cas9 cuts in a mammalian genome, as explored by Canver et al. ^20^, suggested that the frequency of fragment deletion occurring between these cuts is higher at shorter distances. On the contrary, He et al. ^21^ did not observe any apparent correlation between the size of deleted fragment and the deletion frequency. In our experiment, we compared the spacing of two intronic gRNAs designed to target exons of essential genes with the subsequent fold change of the double cut. We observed no significant correlation between the size of target regions, which ranged from 104 to 2132 bp (Fig. 4F). Thus, there does not appear to be an optimum distance between the targeting gRNAs that improves excision efficiency.

To determine whether two flanking gRNAs had a more significant impact compared to the combined effect of individual gRNAs – which may be weak, indirect indication of excision – the genetic interaction score of the gRNA pairs was computed (Fig 5A). We first calculated the expected effect of a pair by summing the singleton effect of each individual gRNA, then computed the delta log fold change (dLFC) as a difference between expected LFC and observed LFC (Fig. 5B). The distribution of dLFC features a median value of 0.09 and a near-symmetric distribution around zero (Fig 5C), suggesting that the simultaneous targeting of flanking sequences both upstream and downstream of essential exons does not exhibit a significant interaction effect compared to targeting each side independently.

**Figure 5.**
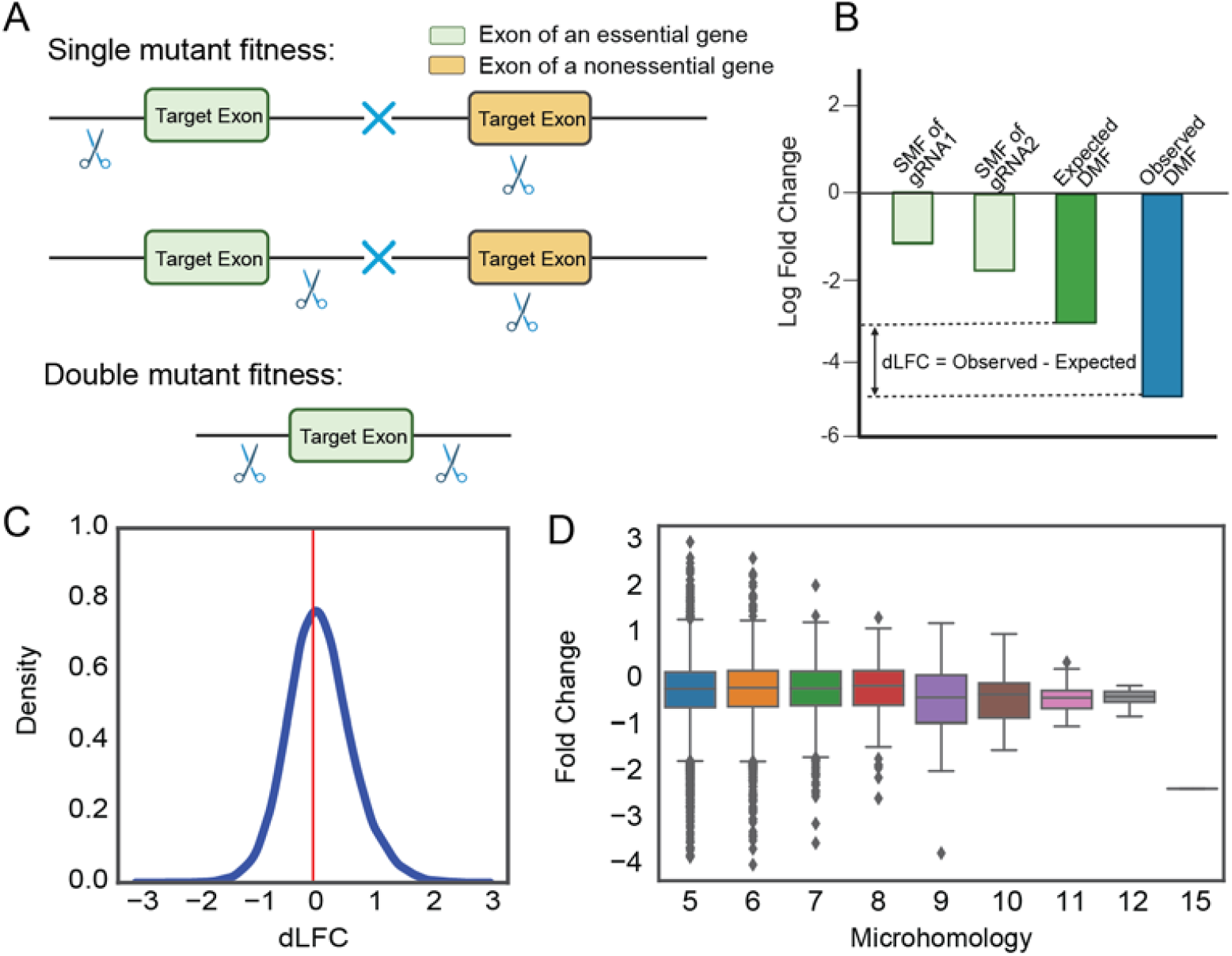
Alternative measures of guide synergy. A) A subset of the library was used to assess the interactions score of intronic gRNAs. B) Quantifying interaction score, dLFC, of intronic gRNAs. Single mutant fitness (SMF) is the mean log fold change of intronic gRNAs. Expected double mutant fitness (DMF) is calculated as the sum of SMF of gRNA1 and gRNA2. Delta Log Fold Change (dLFC) compares observed to expected fold change. C) Density plot of dLFC for intron gRNAs. A left tail is expected for synergistic effects, e.g. locus excision. D) Distribution of fold change across different length of homology between two intronic gRNAs targeting an essential gene.

Finally, we considered the role of DNA repair mechanism. Multiple mechanistic pathways have been identified as efficient mechanisms for repairing double-strand breaks in DNA. The dominant one involves homologous recombination, a process where an undamaged copy of the DNA molecule is used as a template to precisely repair the broken strand. The second pathway is Ku-dependent non-homologous end joining, which directly rejoins the broken ends but can result in some loss of genetic information. A third pathway, microhomology-mediated end joining (MMEJ) operates by utilizing short identical sequences of DNA (microhomologies) near the break sites to guide the repair process. This method often leads to deletions in the repaired DNA sequence, showcasing its distinct mechanism and outcomes compared to the other pathways ^22^. MMEJ relies on these microhomologous sequences spanning 5 to 25 base pairs (bp) on either side of cut sites, and these sequences play a crucial role in aligning the fragmented DNA ends before they are fused together, which ultimately leads to the creation of deletions adjacent to the initial break site ^22^. To investigate the impact of DNA sequence similarity on the excision efficiency in our enCas12a screening, we conducted a comparative analysis of the 30-base sequences flanking each side of the intron cut sites; i.e. comparing upstream of the 5’ cut with downstream of the 3’ cut, to see whether post-excision microhomology improved outcomes. We scored the microhomology of each gRNA by measuring the length of the longest shared sequence between these flanking regions. Microhomology sequences ranging from 5bp to 15bp were observed, but revealed no apparent influence on outcomes (Fig. 5D).

A recent study by Xiao et al. ^23^ employed a similar approach to identify fitness-promoting exons in two cell lines, using the “CHyMErA” method, which co-expresses Cas9 and Cas12a along with libraries of hybrid Cas9/Cas12a guide RNAs. We applied our analysis pipeline to their data and observed patterns similar to those from our data. Positive controls with either one or two guides targeting exons of essential genes showed strong dropout, indicating the CRISPR endonucleases were operating as expected in RPE1 cells (Figure 6A). CHyMErA constructs targeting essential exons did show greater marginal dropout relative to other classes than our Cas12a (Figure 6A, red), but significantly less dropout than constructs directly targeting exons (Figure 6B; Cohen’s D = 1.47). Locus excision CHyMErA constructs did show greater dropout than constructs targeting a single intron near an essential exon (Figure 6C), albeit with much lower magnitude (Cohen’s D=0.69) than between locus excision and direct exon targeting. These findings were repeated in HAP1 cells (Figure 6D-F). Overall, these results are consistent with low-efficiency locus excision using the CHyMErA platform which, though somewhat greater than that we observed with the AsCas12a platform, is still insufficient for negative selection screening.

**Figure 6.**
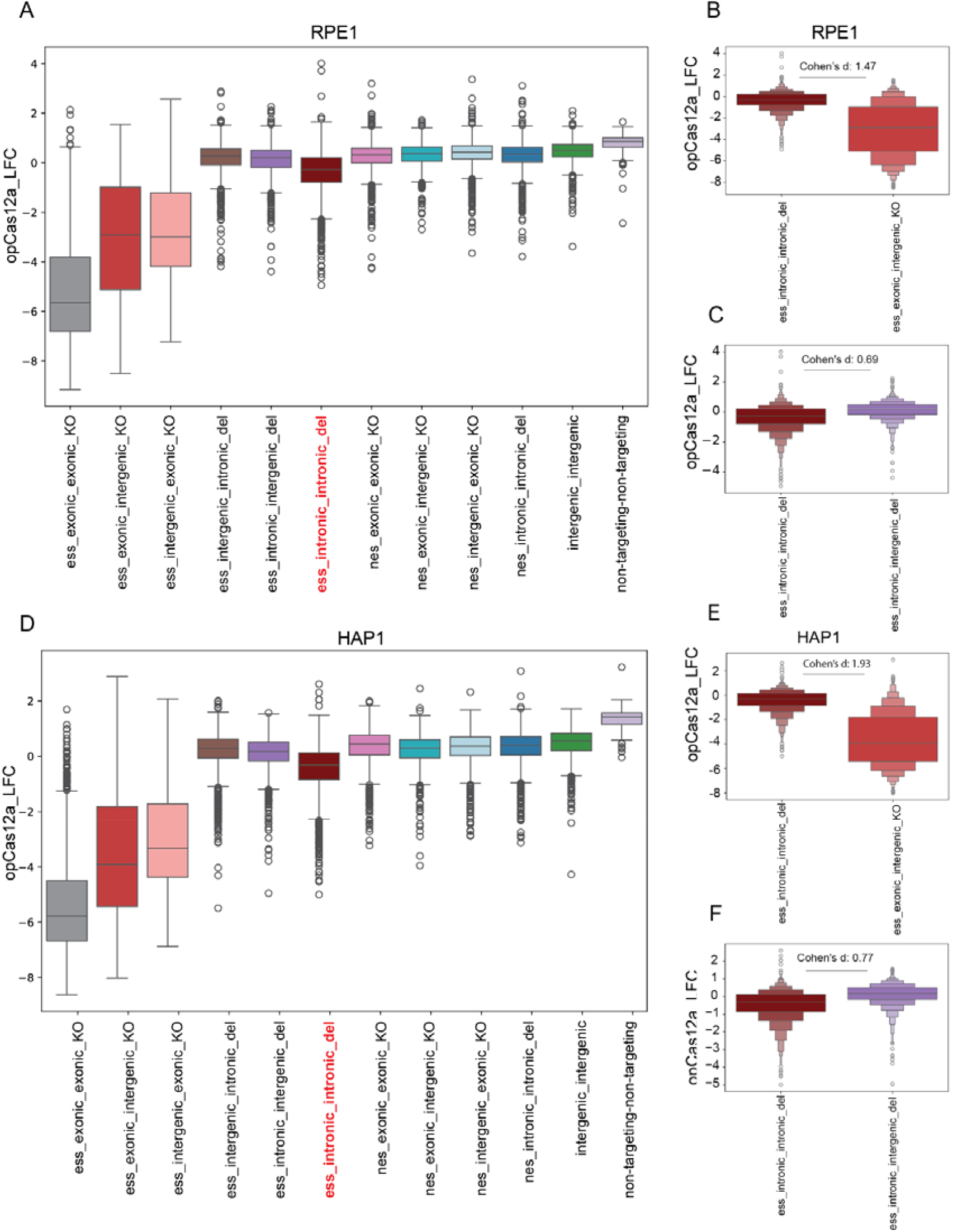
A) Boxplot shows the distribution of fold change in experiment and control groups in Xiao et al.study – RPE1 cell line. B) comparing exonic and intronic arrays targeting essential genes in RPE1 cells. C) comparing locus excision constructs and constructs targeting a single intron near an essential exons in RPE1 cells. D) Boxplot shows the distribution of fold change in experiment and control groups–HAP1 cell line. E) comparing exonic and intronic arrays targeting essential genes in HAP1 cells. F) comparing locus excision constructs and constructs targeting a single intron near an essential exons in HAP1 cells.

## Conclusions

Molecular tools for investigating noncoding regions of the human genome are sparse, an important gap given that less than 2% of the genome consists of protein-coding genes. Recent advancements in CRISPR technology have provided promising avenues for conducting genomic perturbation studies; however, challenges persist when targeting less understood genetic elements or noncoding regions. Excising these regions from the genome could be used as a more potent approach to understand their functional relevance. Our study aimed to explore the capability of Cas12a to cut regions flanking target regions, ideally resulting in their excision from the genome. Our investigation explored identifying high performing gRNA pairs and included a comprehensive exploration of guide and target parameters, such as gRNA orientation, GC content, distance between cuts, and microhomology sequences. While positive controls indicated the targeted endonuclease system was performing as expected, we found no combination of factors that could reliably predict locus excision, and our conclusions are supported by other published data.

Although we do not explore the mechanisms for this failure, our working hypothesis is that either the DNA cuts or the DSB repairs occur asynchronously; logically, both must occur simultaneously for locus excision to result. It is possible that higher concentration of endonuclease could improve the odds of simultaneous DSB, but this is challenging in the context of a lentiviral-mediated pooled screen. We therefore conclude that locus excision, using currently available tools, is not suitable for pooled-library, negative-selection screens, although stringent positive selection experiments could conceivably be viable.

## Supplementary Methods

### Selecting genes

18 essential and 10 nonessential genes were selected from the Hart list of reference genes ^18^ . To reduce any potential bias related to specific chromosomes, genes were distributed across various chromosomes.

### Essential genes

POLR3A, DHX15, TTC27, RPA1, WDR43, SF3B3, LAS1L, NARS1, YARS1, IARS1, KIF23, POLR2B, NOP16, RRM1, INTS9, BRAF, ATP6V1A, CDC16.

### Nonessential genes

TRPM1, ASIC5, TBL1Y, C8B, LIPM, BEND2, SLC36A3, CTCFL, SLC10A2, PGLYRP3.

### Selecting exons

Using gencode.v37, most supported transcript of gene (transcript_support_level = 1) was selected and for each transcript, a selection of 1 to 4 exons was made. We specifically opted for smaller exons, ensuring they were under 200 base pairs in size while still being larger than 35 base pairs to enable the design of exonic gRNAs. The exon sizes were intentionally chosen to be non-multiples of 3 in order to enhance the likelihood of inducing a frameshift. Exons were spaced apart by a minimum of 1100 base pairs to facilitate the design of multiple intronic gRNAs.

### Selecting gRNAs

all the gRNAs were designed using CRISPick. gRNAs targeting selected exons and selected genes (might target different exons) were picked for essential and nonessential genes. Moreover, for the experiment group 1000 bp sequence upstream and downstream of selected exons were used to pick 30 intronic gRNAs for each side. In total, 900 intronic pairs were used to excise each exon.

For generating construct to target exons in essential and nonessential genes, we made construct that the first guide RNA was an exonic gRNA targeting one of the picked exons in the target genes and the second guide RNA was a random gRNA from our pool of negative control guides that they either target nonessential genes or are non-targeting guides.

To generate the construct to excise selected loci we selected the intronic guides downstream or upstream of the picked exons.

#### Cell Culture

A375 cells were obtained from the Cancer Cell Line Encyclopedia (Broad Institute) and grown in RPMI+10% FBS. Cells lines were routinely tested for Mycoplasma contamination and were maintained without antibiotics except during screens, when the medium was supplemented with 1% penicillin–streptomycin. Cell lines were kept in a 37 °C humidity-controlled incubator with 5.0% carbon dioxide and were maintained in exponential phase growth by passaging every 2 – 4 days.

#### enCas12a Screen

Cells were first engineered to express EnAsCas12a using the vector pRDA_174, described previously. Cells were then transduced in 3 biological replicates with a lentiviral library (CP1695). Transductions were performed at a low multiplicity of infection (MOI ∼0.35), using enough cells to achieve a representation of at least 500 transduced cells per construct. Puromycin (1ug/mL) was added post-transduction to remove non-transduced cells. The culture was passaged every 2 - 3 days for 21 days. Cells were pelleted by centrifugation, resuspended in PBS, and frozen promptly for genomic DNA isolation.

### Genomic DNA preparation and sequencing

Genomic DNA (gDNA) was isolated using the KingFisher Flex Purification System with the Mag-Bind Blood & Tissue DNA HDQ Kit (Omega Bio-Tek. The gDNA concentrations were measured by Qubit. For PCR amplification, gDNA was divided into 100 µL reactions such that each well had at most 10 µg of gDNA. Plasmid DNA (pDNA) was also included at a maximum of 100 pg per well. Each well of a 96-well PCR plate contained 1.5 µL of Titanium Taq (Takara), 10 µL of Titanium Taq buffer, 8 µL of dNTPs, 5 µL of DMSO, 0.5 µL of P5 primer at 100 µM stock, 10 µL of P7 primer at 5µM stock, and sterile water added to 100 µL. PCR cycling conditions were as follows: (1) 95°C for 1 min; (2) 94°C for 30 s, (3) 52°C for 30 s, (4) 72°C for 30 s, (5) go to step 1, x28; (6) 72°C for 10 min. PCR products were purified with Agencourt AMPure XP SPRI beads according to manufacturer’s instructions (Beckman Coulter, A63880). Samples were sequenced via the Broad Genomics Platform Walk-Up Illumina sequencing service.

#### QC analysis

The subsequent analysis was performed using Python notebooks (Python version: 3.10.12.). The mean read depth for all samples was greater than 500 reads per guide. A pseudocount of 5 reads was added to each array in every sample, and read counts were normalized to an average of 500 reads per guide to assess the correlation between different samples and replicates.

#### Cohen’s D

Cohen’s d between the fold change distributions of different groups of guide arrays, as well as between guide array orientations, was calculated using NumPy package as follows:

Where:

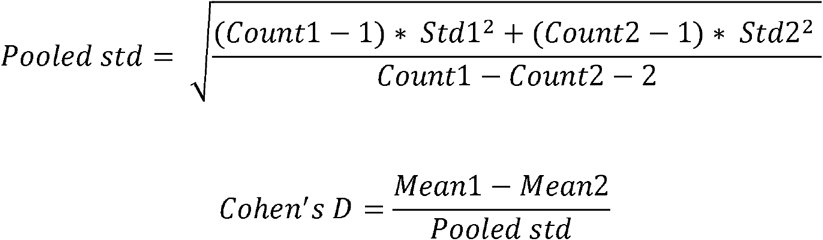

Mean1 = Mean LFC of arrays targeting group 1

Mean2 = Mean LFC of arrays targeting group 2

Std1 = Standard deviation of LFC of arrays targeting group 1

Std2 = Standard deviation of LFC of arrays targeting group 2

Count 1 = Number of arrays targeting group 1

Count 2 = Number of arrays targeting group 2

#### Double knockout phenotype

For target exons in essential genes, effect of each intronic gRNA, single intron mutant fitness (SMF), was calculated as the mean construct log fold change of intronic-control constructs. The control was either non-essential genes or safe-targeting gRNAs. For each exon, the expected double mutant fitness (DMF) of intronic gRNA 1 and intronic gRNA 2 was calculated as the sum of SMF1 and SMF2. The difference between expected and observed DMF was called dLFC.

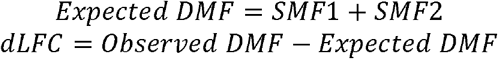

#### Microhomology

A microhomology score was determined by comparing the 30-base DNA sequence before upstream gRNA cuts with the 30-base DNA sequence after downstream cuts, identifying the length of the longest matching sequence between them. This measurement quantifies the extent of homology between the two DNA segments.

### Calculation of Coefficient of Determination (R^2^)

We used simple linear regression to assess the relationship between two variables. A line was fitted using numpy. polyfit, and the R^2^ value was calculated with the r2_score function from scikit-learn to show how well the line explained the data.

## Reference

1. Cheng, Y. et al. Principles of regulatory information conservation between mouse and human. Nature 515, 371–375 (2014).

2. Maurano, M. T. et al. Systematic Localization of Common Disease-Associated Variation in Regulatory DNA. Science 337, 1190–1195 (2012).

3. Montalbano, A., Canver, M. C. & Sanjana, N. E. High-Throughput Approaches to Pinpoint Function within the Noncoding Genome. Molecular Cell 68, 44–59 (2017).

4. Shukla, A. & Huangfu, D. Decoding the noncoding genome via large-scale CRISPR screens. Current Opinion in Genetics & Development 52, 70–76 (2018).

5. Diao, Y. et al. A tiling-deletion-based genetic screen for cis-regulatory element identification in mammalian cells. Nat Methods 14, 629–635 (2017).

6. Aparicio-Prat, E. et al. DECKO: Single-oligo, dual-CRISPR deletion of genomic elements including long non-coding RNAs. BMC Genomics 16, 846 (2015).

7. Gasperini, M. et al. CRISPR/Cas9-Mediated Scanning for Regulatory Elements Required for HPRT1 Expression via Thousands of Large, Programmed Genomic Deletions. The American Journal of Human Genetics 101, 192–205 (2017).

8. DeWeirdt, P. C. et al. Optimization of AsCas12a for combinatorial genetic screens in human cells. Nat Biotechnol 39, 94–104 (2021).

9. Zetsche, B. et al. Cpf1 Is a Single RNA-Guided Endonuclease of a Class 2 CRISPR-Cas System. Cell 163, 759–771 (2015).

10. Parrish, P. C. R. et al. Discovery of synthetic lethal and tumor suppressor paralog pairs in the human genome. Cell Rep 36, 109597 (2021).

11. Thompson, N. A. et al. Combinatorial CRISPR screen identifies fitness effects of gene paralogues. Nat Commun 12, 1302 (2021).

12. Ito, T. et al. Paralog knockout profiling identifies DUSP4 and DUSP6 as a digenic dependence in MAPK pathway-driven cancers. Nat Genet 53, 1664–1672 (2021).

13. Gonatopoulos-Pournatzis, T. et al. Genetic interaction mapping and exon-resolution functional genomics with a hybrid Cas9–Cas12a platform. Nat Biotechnol 38, 638–648 (2020).

14. Kleinstiver, B. P. et al. Engineered CRISPR–Cas12a variants with increased activities and improved targeting ranges for gene, epigenetic and base editing. Nat Biotechnol 37, 276–282 (2019).

15. Esmaeili Anvar, N. et al. Efficient gene knockout and genetic interaction screening using the in4mer CRISPR/Cas12a multiplex knockout platform. Nat Commun 15, 3577 (2024).

16. Lenoir, W. F. et al. Discovery of putative tumor suppressors from CRISPR screens reveals rewired lipid metabolism in acute myeloid leukemia cells. Nat Commun 12, 6506 (2021).

17. Dede, M., McLaughlin, M., Kim, E. & Hart, T. Multiplex enCas12a screens detect functional buffering among paralogs otherwise masked in monogenic Cas9 knockout screens. Genome Biology 21, 262 (2020).

18. Hart, T., Brown, K. R., Sircoulomb, F., Rottapel, R. & Moffat, J. Measuring error rates in genomic perturbation screens: gold standards for human functional genomics. Mol Syst Biol 10, 733 (2014).

19. Hart, T. et al. High-Resolution CRISPR Screens Reveal Fitness Genes and Genotype-Specific Cancer Liabilities. Cell 163, 1515–1526 (2015).

20. Canver, M. C. et al. BCL11A enhancer dissection by Cas9-mediated in situ saturating mutagenesis. Nature 527, 192–197 (2015).

21. He, Z. et al. Highly efficient targeted chromosome deletions using CRISPR/Cas9. Biotech & Bioengineering 112, 1060–1064 (2015).

22. McVey, M. & Lee, S. E. MMEJ repair of double-strand breaks (director’s cut): deleted sequences and alternative endings. Trends in Genetics 24, 529–538 (2008).

23. Xiao, M.-S. et al. Genome-scale exon perturbation screens uncover exons critical for cell fitness. Molecular Cell 84, 2553-2572.e19 (2024).

